# A structural classification of candidate oscillators and multistationary systems

**DOI:** 10.1101/000562

**Authors:** Franco Blanchini, Elisa Franco, Giulia Giordano

**Affiliations:** Dipartimento di Matematica ed Informatica, Università degli Studi di Udine, Via delle Scienze 206, 33100 Udine, Italy; Department of Mechanical Engineering, University of California, Riverside, 900 University Avenue, Riverside, CA 92521

**Keywords:** Feedback loops, multistationarity, stability, oscillator, structural properties

## Abstract

Molecular systems are uncertain: the variability of reaction parameters and the presence of unknown interactions can weaken the predictive capacity of solid mathematical models. However, strong conclusions on the admissible dynamic behaviors of a model can often be achieved without detailed knowledge of its specific parameters. In particular, starting with Thomas' conjectures, loop–based criteria have been largely used to characterize oscillatory and multistationary dynamic outcomes in systems with a sign definite Jacobian.

We build on the rich literature focused on the identification of potential oscillatory and multistationary behaviors based on parameter–free criteria. We propose a classification for sign–definite non autocatalytic biological networks which summarize several existing results in the literature, adding new results when necessary. We define candidate oscillators and multistationary systems based on their admissible transitions to instability. We introduce four categories: strong/weak candidate oscillatory/multistationary systems, which correspond to networks in which all/some of the existing feedback loops are negative/positive. We provide necessary and sufficient conditions characterizing strong and weak candidate oscillators and multistationary systems based on the exclusive or simultaneous presence of positive and negative loops in their linearized dynamics. We also consider the case in which the overall system is the connection of several stable aggregate monotone components, providing conditions in terms of positive/negative loops in a suitable network with aggregate monotone systems as nodes.

Most realistic examples of biological networks fall in the gray area of systems in which both positive and negative cycles are present: therefore, both oscillatory and bistable behavior are in principle possible. Native systems with a large number of components are often interconnections of monotone modules, where negative/positive loops among modules characterize oscillatory and bistable behaviors, in agreement with our results. Finally, we note that many canonical example circuits exhibiting oscillations or bistability fall in the categories of strong candidate oscillators/multistationary systems.

## 1 Introduction

What interaction networks can generate multistationary and periodic behaviors in molecular systems? This is a fundamental question in the area of systems and synthetic biology, as it is relevant both when *analyzing* native biological circuits, and when *building* artificial networks to achieve target dynamics. However, the variability and uncertainty characterizing molecular circuits pose significant challenges to answering such question. Mathematical models facilitate and support the analysis and interpretation of uncertain dynamic data: but models are often plagued by uncertainty as well, whenever it is impractical (if not impossible) to measure their parameters. Thus, from a modeling perspective, satisfactory answers to our question may be found by identifying “structural” sources of multistationarity and periodicity, seeking parameter–free criteria to predict the capacity of a model to exhibit specific dynamic “phenotypes”.

Parameter–free criteria to determine local admissible dynamics of a system can be formulated by studying Jacobian graphs. Concentrations of molecular species (states of the dynamical system) are associated with nodes of the graph, and signed Jacobian entries linking different states are associated with directed, signed edges of the graph. Checking for the presence of positive or negative loops in the Jacobian graph is a widely accepted method to explain multistationarity and oscillations in molecular/chemical systems [51]. Some of the first and best known mathematical conjectures in this area were formulated by R. Thomas [49]: given a Jacobian graph, a negative loop is a necessary condition for stable periodic behavior, while a positive loop is a necessary condition for multistationarity (see [13] for a very thorough survey). These conjectures were proved in [22] and [45], with several further extensions and refinements [7, 27, 42, 47].

While Thomas' loop conditions are only necessary (the dynamic phenotype of a system always depends on its specific parameters), they provide useful guidelines for analysis and design. In particular, design of feedback loops underlies the successful creation of many novel artificial biochemical networks capable of bistability [6, 20, 29, 40] or oscillations [16, 19, 28, 37, 48, 50]. In practice, however, it is extremely challenging to generate “clean” feedback loops in any experimental system. Even *in vitro* artificial networks [19, 28, 37] with a small number of components are generally affected by unknown and undesired interactions among reactants, which can create additional parasitic feedback loops of uncertain strength. This challenge motivates the search for structural criteria to discriminate dynamic outcomes of a network where several potentially unknown loops coexist.

We contribute to the research in this direction by proposing a structural classification where we contrast *weak* and *strong candidate* oscillators and multistationary systems; our classification depends on the exclusive or simultaneous presence of positive and negative loops. We consider non–autocatalytic systems having a sign definite Jacobian, and we identify the *structure* of a system with the sign pattern of its Jacobian and the corresponding directed graph. A structure can be specified into a dynamic *realization* by choosing a set of model parameters. Our definition of candidate dynamic behaviors is based on the admissible types of *transition to instability*, which occurs when the eigenvalues of the Jacobian cross the imaginary axis, due to a change in parameters. Multistationarity is associated with real eigenvalues transitioning to instability, while oscillatory behaviors are associated with pairs of complex eigenvalues transitioning to instability. It is well known that these phenomena are related to bifurcation theory. Typically, a transition on the real axis is related to the presence of a pitchfork bifurcation, while a transition with complex eigenvalues is related to a Hopf bifurcation. Formally, however, these types of bifurcation occur under a range of additional assumptions which complicate their mathematical analysis; see, for instance, [2, 34].

Our classification work builds on a rich body of literature [7, 22, 27, 42, 45, 47, 49]. Our proofs rely on the so–called degree theory [24, 39] and on the properties of sign–definite systems [33]. We extend our results to aggregate monotone systems, which are defined as the interconnection of several input–to–state monotonic subsystems. Our approach can be used to characterize systems affected by delays, which are often present in models of gene networks [31]. While the analysis of networks with delays is outside of the scope of this paper, in Appendix D we propose loop–based sufficient conditions to identify candidate oscillators and multistationary systems when delays are present.

### 1.1 Motivating example

We begin with a simple example to introduce some mathematical notions used throughout this paper, and to clarify the relevance of our study in the context of biomolecular systems.

We consider a standard model for transcription and translation of two genes, where proteins reciprocally modulate their expression forming a feedback loop. Similar models are commonly encountered in the literature; see, for instance [16, 28]. For illustrative purposes, we use a nondimensional model:

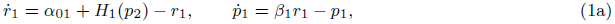

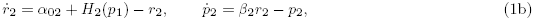

where, for *i* = 1, 2, *r_i_* are RNA species concentrations; *p_i_* are protein concentrations; *H_i_*(·) are Hill functions, and all Greek letters denote reaction rates that are positive scalars. Nondimensionalization is justified and carried out in detail in Appendix A.

Depending on the regulatory action of the proteins, and thus depending on the type of Hill function, the network presents a different number of equilibria and possible dynamic behaviors. For example, suppose 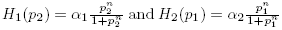: this is a two–gene positive feedback loop, which is often encountered in developmental networks [1, 12]. The Jacobian of the system is:

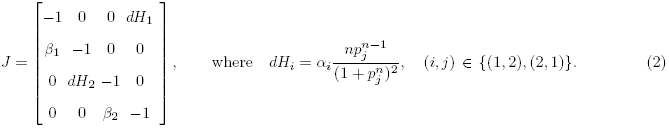

The first remarkable feature is that the Jacobian entries are sign definite, i.e. they do not change sign for arbitrary choices of the (positive) parameters *α_i_, β_i_* and *n*.

The Jacobian sign pattern is a “structural” property of this system, and it can be associated with a graph: nodes correspond to the biological species, and are interconnected by positive or negative arcs according to the corresponding Jacobian entries, as shown in Figure 1 a. We also remark that the system is dissipative, i.e. there is spontaneous degradation of each species, and the diagonal terms of the Jacobian are negative definite. Thus, every node has a negative self–loop.

We can derive expressions for the equilibria of the system, which are given by the intersections of the two nullclines (Figure 1 a, top row):

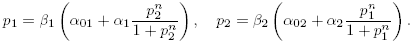

For *n* = 1 we have an additional intersection with *p_1_* and *p_2_* positive; for *n* > 1, the system admits additional (at most two) positive equilibria. Increasing *n* not only introduces new equilibria, but also influences their nature. We can see this by finding the eigenvalues of the Jacobian, which are the roots of its characteristic polynomial:

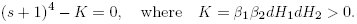

**Fig. 1.**
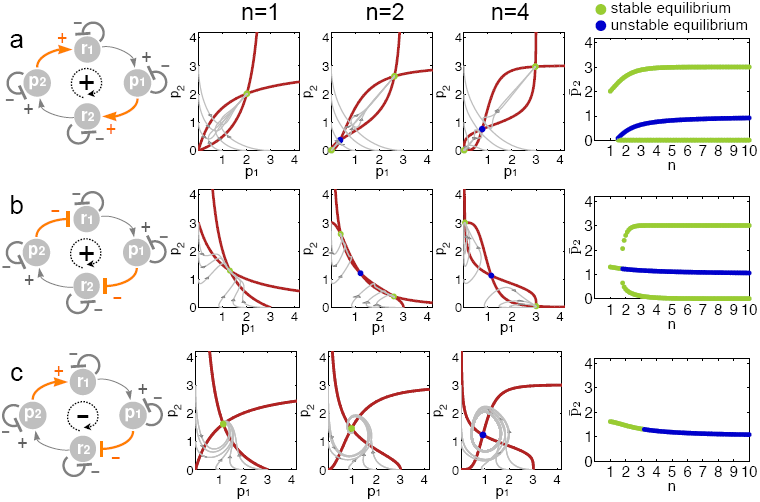
a: Two–gene system with double positive feedback loop. Pointed arrowheads indicate positive Jacobian interconnection entries, while hammer–arrowheads indicate negative interconnections. b: Two–gene system with double negative feedback loop, resulting in an overall positive feedback interconnection. c: Two–gene feedback interconnection with positive and *n* negative regulation, resulting in an overall negative loop. The nondimensional parameters are chosen as *α*_01_ = *α*_02_ = 0.01, *α*_1_ = *α*_2_ = 3, *β*_1_ = *β*_2_ = 1 and *n* is varied. The right column shows the corresponding value of *P*_2_ equilibria for varying, *n* and their different pattern of transition to instability. Green dots are stable equilibria, blue dots are unstable equilibria.

Note that *K* depends on the partial derivatives of the Hill functions, and thus on *n.* If *K* > 1 (i.e., if the Hill coefficients are suitably high) only one of the roots of this polynomial has positive real part, therefore the root must be on the real axis. If *K* < 1, all roots have negative real part. Thus, the only type of instability admitted by the system is real exponential. New equilibria appear as the system transitions to instability. Figure 1 a, top row, shows nullclines, example trajectories in the *p*_1_ – *p*_2_ plane of the phase space, and the equilibrium values of *p*_2_ for varying *n* (stable equilibria are represented as green circles, unstable equilibria as blue circles).

If 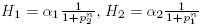, network (1) specifies a two–gene double negative feedback loop, depicted in Figure 1 b, left. This circuit is also known as toggle switch, an example of which is the famous synthetic biological circuit by Gardner [20]; a natural example of a toggle switch is the Cdc2–Wee1 network considered, for instance, in [3]. We can repeat the same analysis done for the two–gene double positive loop, and get similar results in terms of admissible transitions to instability, which can be only real exponential, regardless of the equilibrium considered (Figure 1 b, right panel).

We now compare the previous two examples to the case when Hill functions have opposite regulatory roles, i.e. 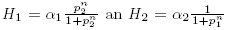: the network can behave as a two–gene oscillator [28]. First, we observe that the Jacobian is still a sign definite matrix; however, the “interconnection” terms *dH*_1_ and *dH*_2_, the derivatives of the Hill functions, now have opposite signs due to the different slopes of such functions, and thus generate an overall *negative* feedback loop (Figure 1 c). In addition, the nullclines are now:

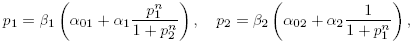

and admit a single intersection regardless of the value of *α*_*i*_, **β**_*i*_, and *n* (Figure 1 c, central panels show the nullclines for increasing values of *n*). The characteristic polynomial is the following:

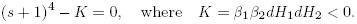

Since now *K* < 0 (because *dH*_2_ is negative, while *dH*_1_ is positive), all the coefficients of this polynomial are positive: therefore, roots with a positive real part are necessarily complex. As a consequence, only oscillatory unstable dynamics can arise, rather than real exponential. A transition to instability does not result in new equilibria.

To summarize, in this simple two–gene system we can reach very strong conclusions regarding its possible dynamic behaviors. Such conclusions do not depend on specific parameter choices, but rather on the positive or negative feedback interconnection. In particular, this example highlights clearly that there is a relationship between feedback loops and admissible transitions to instability. A qualitatively similar study was carried out and validated by building synthetic bacterial circuits in [6]; analysis relied on the S–systems formalism [43].

In the next sections, we propose a general framework to categorize oscillatory and multistationary systems based on the exclusive or coexisting presence of positive and negative feedback loops.

## 2 Assumptions and definitions

We consider dynamical systems:

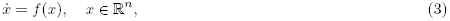

where *f*(·) is continuously differentiable in all its components *f*_*i*_(·), *i* = 1,…, *n*, and may be an uncertain function (for example, Hill coefficients or degradation rates may be fitted from noisy data).

We assume that all the solutions of system (3) are globally uniformly asymptotically bounded in a ball *𝒮*: formally, we say that for any compact set *𝒮*_0_ including *𝒮*, there exists *T* > 0 (depending on *𝒮*_0_) such that *x*(*t*) ∈ *𝒮* for any *x(0)* ∈ *𝒮*_0_ and any *t* ≥ *T*. As a consequence, the system admits an equilibrium point *x̄* in *𝒮*, and without restriction we may assume that *x̄* = 0.

### Assumption 1

*We assume that each component f_i_(·) is monotone (either non–increasing or non–decreasing) with respect to each argument x_i_*.

This is equivalent to assuming that the Jacobian of the system is sign definite, i.e. *∂f_i_/∂x_j_* is either always positive, always negative, or always null in all the considered domain.

### Assumption 2

*We assume that the system is locally dissipative, i.e. ∂f_i_/∂x_i_ < 0. Equivalently, we say that the systems considered are non–autocatalytic.*

Because of the monotonicity assumption, a sign pattern matrix *Σ* can be immediately associated with the sign definite Jacobian of system (3). Zero elements in the Jacobian correspond to zero elements in *Σ.* We refer to *Σ* as the *system structure.* This structure is associated with a directed, *n*-node graph, whose arcs are positive, negative, or zero depending on the sign of the corresponding entries in *Σ*. We provided several graph examples within the two–gene network considered in Figure 1.

Given a system structure *Σ*, we call a *realization* of such structure any choice of functions *f*_*i*_(·), and thus of the Jacobian entries *J* = [*∂f_i_/∂x_j_*].

### Definition 1

We say a property *𝒫* is structural if it is satisfied by a system structure, independent of the specific realization [8].

For example, the structure corresponding to the two–node double positive feedback loop is:

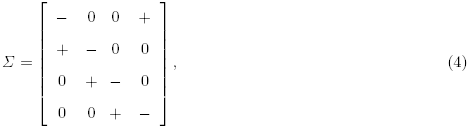

as shown in Figure 2. The specific model we considered and its Jacobian matrix (2) are a possible realization of this structure.

### Definition 2

A cycle is an oriented, closed sequence of distinct nodes connected by distinct directed arcs in the graph of a structure. We say the loop is negative if the number of negative arcs is odd; otherwise, we say the loop is positive. The number of arcs forming a loop is called the order of the loop.

**Fig. 2.**
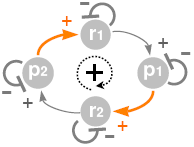
Graph corresponding to the structure at Equation 4.

Referring to our two–genes example network, all three cases have a single loop. However, the double positive feedback system and the toggle switch exhibit an overall positive loop, while the two–node oscillator presents a single negative loop.

The order of the cycles in a system can be an important factor in determining its dynamic behavior. Systems where all the negative cycles (if any) are of order two present particular challenges: we denote these systems as critical, and discuss the validity of our results for this class of structures in Appendix C. In the remainder of this paper, we consider non–critical systems.

We now introduce general definitions of transition to instability. Consider the parameter–dependent dynamical system:

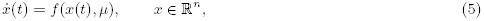

where *f*(·, ·) is a sufficiently smooth function, and μ is a real–valued parameter. Recalling our boundedness assumption, we assume that an equilibrium exists and it is the origin, i.e. *f*(0,*μ*) = 0. In general, any equilibrium point is a function of parameter *f*(*x̄_μ_,μ*) = 0; however, a suitable change of coordinates always allows us to shift the equilibrium to the origin, without affecting our analysis.

### Definition 3 Transition to Instability (TI)

System (5) undergoes a transition to instability for *μ = μ\** iff its Jacobian matrix *J* is an asymptotically stable matrix in a left neighborhood of *μ*\*, and it is an unstable matrix in a right neighborhood^1^.

In the paper we limit our attention to *simple* TIs, in the sense that at most a single real eigenvalue or a pair of complex conjugate eigenvalues crosses the imaginary axis. Note that, in principle, many eigenvalues could leave the stability region simultaneously. However, this is not likely to occur in realistic situations. In addition, in most systems one can identify a *dominant* eigenvalue, *i.e*. an eigenvalue (or a pair of complex conjugate eigenvalues) having real part larger than any other eigenvalue; a dominant eigenvalue is the primary parameter defining the dynamic response of a system. Here, we assume that the eigenvalues generating a TI are dominant.

We now specify two types of TIs that are related to oscillatory and multistationary dynamic behaviors.

### Definition 4 Oscillatory Transition to Instability (OTI)

System (5) undergoes a (simple) oscillatory transition to instability for *μ* = *μ\** iff its Jacobian matrix *J* has a simple pair of pure imaginary eigenvalues, while all the other eigenvalues have negative real part:

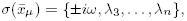

with *Re(λ_k_)* < 0 for *k* ≥ 3 and with *Re(λ_k_)* > 0, for *k* = 1, 2, in a right neighborood of *μ\**.

### Definition 5 Monotonic Transition to Instability (MTI)

System (5) undergoes a (simple) monotonic or real exponential transition to instability for *μ* = *μ\** iff the Jacobian matrix *J* has a zero simple eigenvalue, while all remaining eigenvalues have negative real part:

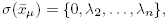

with the condition *Re(λ_k_)* < 0 for *k* ≥ 2 and with *Re(λ_1_)* > 0, in a right neighborhood of *μ*^*^.

### Definition 6

We say that the transition to instability is *simple* if it is either a OTI or a MTI.

When an MTI occurs, the determinant of the Jacobian changes sign as the parameter crosses the stability limit. Therefore, under overall boundedness assumptions, an MTI causes the appearance of new equilibrium points [34]. This is not the case for OTIs. OTIs are generally related to Andronov–Hopf bifurcations, while MTIs are associated with pitchfork bifurcations. However, these types of bifurcation can only occur under additional non–singularity assumptions; for example, it is required that the derivative of the dominant eigenvalues with respect to the changing parameter be strictly positive [2, 34].

## 3 Structural classification

We begin by stating general definitions for candidate oscillatory and multistationary networks. Based on these definitions, we then propose necessary and sufficient conditions that link the presence of positive and negative loops in a structure to the oscillatory or multistationary nature of the system. A table summarizing our classification based on loops in the system structure is in Figure 3.

### Definition 7

System (3) is structurally a candidate oscillator in the weak sense iff, for some realization of the considered functions, it admits an OTI.

**Fig. 3.**
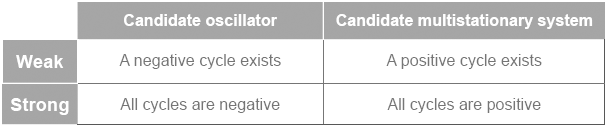
Table summarizing our structural loop–based classification of candidate oscillators and multistationary systems.

### Definition 8

System (3) is structurally a candidate oscillator in the strong sense iff every simple transition to instability (if any) is an OTI.

As a “dual case” we consider the case of candidate multistationary systems.

### Definition 9

System (3) is structurally a candidate multistationary system in the weak sense iff, for some realization of the considered functions, it admits a simple MTI.

### Definition 10

System (3) is structurally a candidate multistationary system in the strong sense iff every simple transition to instability (if any) is an MTI.

For example: a weak candidate oscillator admits, for some choice of the parameters, an unstable equilibrium point where trajectories spiral out of the equilibrium with oscillatory dynamics. A strong candidate oscillator structure admits transitions to instability exclusively of oscillatory nature. The two–gene oscillator system described in the previous section is a strong candidate oscillator. In fact, unstable dynamics are always associated with complex conjugate eigenvalues, and are therefore oscillatory. Conversely, a weak candidate multistationary structure admits, for some choice of the parameters, monotonically unstable behaviors. A strong candidate multistationary system admits solely monotonic unstable dynamics. Going back to our examples, both the two–gene double positive loop and the toggle switch are strong candidate multistationary systems. Unstable equilibria can only have monotonic nature.

We are now ready to state our main results. The following propositions provide necessary and sufficient conditions for candidate oscillators and multistationary systems, based on the sign of the loops present in their structure. The complete demonstration of each proposition is reported in the Appendices; here we only sketch the main idea of each proof.

### Proposition 1

*A non–critical system is a candidate oscillator in the weak sense if and only if its structure has at least one negative cycle (necessarily of order greater than 2)*.

The necessity part of the proof is based on the well known fact that systems with only positive cycles are monotone, thus cannot admit undamped oscillations [44, 46]. The sufficiency part of the proof relies on the application of a special class of transformations (see Appendix B.1) allowing to independently scale the magnitude of desired loops in a realization. The complete proof is in Appendix B.2. Weak candidate oscillators are exemplified in the literature by the amplified negative feedback oscillators and incoherent oscillators described in [38] and by the toggle–switch/oscillator circuit in [6].

### Proposition 2

*A non–critical system is a candidate oscillator in the strong sense if and only if its structure has only negative cycles*.

The necessity part of this proposition can be demonstrated similarly to the sufficiency part of Proposition 1. The sufficiency part can be demonstrated using algebraic tools from [33]: a structure with only negative loops can only present complex unstable eigenvalues (OTI). See Appendix B.3 for the complete proof. Examples of strong oscillators are the well known repressilator circuit [16], and all the negative feedback oscillator models described in [38].

### Corollary 1

*A strong candidate oscillator admits a single equilibrium*.

This corollary is a consequence of degree theory: see Appendix B.4.

### Proposition 3

*A non–critical system is a candidate multistationary system in the weak sense if and only if its structure has at least a positive cycle*.

The necessity part of this proof is done by contradiction: if no positive loop is present, then the system is a candidate oscillator in the strong sense. Sufficiency is demonstrated by introducing transformations that allow independent scaling of Jacobian entries in a realization; this is the same argument given in the necessity part of the proof of Proposition 2. The complete proof is reported in Appendix B.5. Examples of candidate weak multistationary systems include the amplified negative feedback oscillators and incoherent oscillators in [38], and again the toggle–switch/oscillator circuit in [6].

### Proposition 4

*A non–critical system is a candidate multistationary system in the strong sense if and only if its structure has positive cycles only*.

The necessity part of this proposition is easily proved by contradiction: if a negative cycle exists, then one can find a realization exhibiting an OTI, according to Proposition 1. Sufficiency follows due to the fact that a structure with only positive cycles gives rise to a monotone system; thus, only MTI is admitted. The complete demonstration can be found in Appendix B.6.

Examples of strong candidate multistationary systems include the previously presented two–gene double positive loop and toggle switch architectures [3, 20].

### Corollary 2

*A strong candidate multistationary system, in which a simple MTI occurs at μ*, admits additional stationary points in a right (unstable) neighborhood of μ\**.

*If these additional equilibria are non–singular (i.e. if the determinant of the Jacobian is non–zero), then there are at least two additional equilibria*.

*If exactly two additional non–singular equilibria appear, and the Jacobian evaluated at those points, J(x̄), has a single dominant eigenvalue (i.e. the system is irreducible), then these equilibria are asymptotically stable*.

The proof is reported in Appendix B.7.

*Remark 1* Delay differential equations are often adopted to model molecular systems. For example, transcription, processing and transport of mRNA in genetic networks has been successfully modeled with explicit delays; see, for instance, [31]. A thorough analysis of oscillations and multistationarity in delay differential equations is out of the scope of this paper. However, we discuss this challenge in Appendix D and we provide local, loop–based sufficient conditions for OTIs and MTIs in a wide class of systems affected by delays.

### 3.1 Oscillations and multistationarity in aggregates of stable monotonic systems

Many biological networks fall into the category of monotone systems [46]. Monotonicity is a property that can be verified without exact knowledge of the system parameters; thus, criteria relying on monotonicity can be in turn considered robust with respect to parametric uncertainty. We extend our results to “aggregate” systems that are composed of monotonic subsystems, for which we provide definitions below. Many biological systems have been analyzed, from a systems perspective, as the interconnection of monotone subsystems; for example, the Cds–Wee1 network [3], the MAPK pathway [46], and the Goldbeter oscillator [4, 21] in *Drosophila*. An aggregate system is the interconnection of *N* subsystems defined as follows:

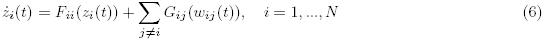

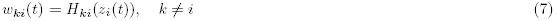

where *z_i_*(*t*) is a vector associated with the *i*-th subsystem. Subsystem *i* receives inputs from subsystems *j*, and sends an output to subsystem *k* (see Figure 4). Functions *G_ij_*(*w_j_*(*t*)) model the influence of subsystem *j* on subsystem *i*, through the nonnegative real variable *w_ij_*(*t*), output of subsystem *j*.

**Fig. 4.**
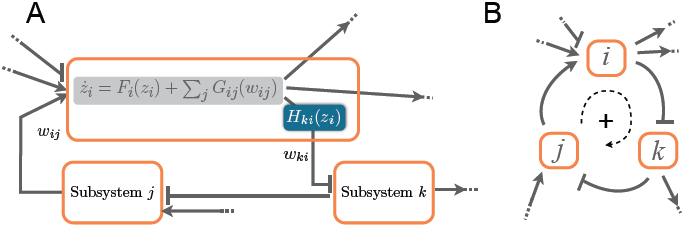
A: Sketch of an aggregate monotone system, which is the interconnection of several monotone systems, defined in equations (6)–(7); pointed arrowheads indicate non–decreasing interconnections, and hammer–arrowheads indicate non– increasing interconnections. B: Graph of the aggregate system where each monotone subsystem is collapsed in a single node.

#### Assumption 3

*We assume that functions G_ij_(w_ij_) are non–decreasing*.

The above assumption is not restrictive and enables a simplified analysis. “Negative” interconnection trends among subsystems can be captured by the output functions *w_ij_ = H_ij_(z_i_)*. For example, consider a generic node 1, and the influence of node 2 on 1 given by *w*_12_:

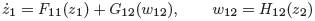

with *G*_12_ decreasing. The overall interaction depends on the monotonic compound function *G*_12_ ° *H*_12_. Thus, we can “move” the decreasing interaction trend from *G*_12_ to *H*_12_ with a simple sign change: *ŵ*_12_ = −*w*_12_. The overall result of the compound function remains unchanged:

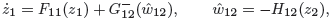

where 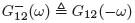 is now increasing.

#### Assumption 4

*We assume that the state–to–output mapping of each subsystem is either non–decreasing or non–increasing: i.e. 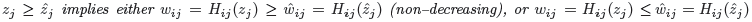*.

If an input/output monotonicity characterization is available for each subsystem, we will show that: 1) We can collapse each subsystem into an equivalent node, and 2) We can still classify structural oscillatory and multistationary behaviors based on the loops created by the interconnections among these new, equivalent nodes.

#### Definition 11

The *i*–th subsystem is unconditionally stable iff, for constant input values *w̄*_ij_, it admits a single equilibrium *z̄*_*i*_, the solution of:

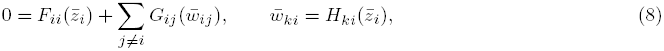

and the equilibrium is globally stable (all the eigenvalues of the *i*–th Jacobian have a negative real part).

We apply the usual definition of input–to–state monotonicity.

#### Definition 12

We say that the i–th subsystem (6)-(7) is input–to–state monotone iff it has the following property:
 

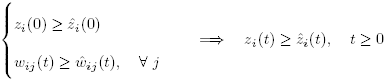

#### Assumption 5

*We consider aggregate systems where each subsystem satisfies definitions 11 and 12*.

We can now define an **aggregate graph** (Figure 4 B): each subsystem is collapsed into a node, and any interconnection between the *i*–th and *k*–th subsystem is a directed arc with a sign that depends on the trend of the corresponding state–to–output mapping *H_ki_(z_i_)*. (Positive arcs are associated with non–decreasing mappings, negative arcs are associated with non–increasing mappings.)

#### Proposition 5

*Consider an aggregate system, formed by the interconnection of several monotone unconditionally stable subsystems as* (6)–(7), *satisfying Assumptions 3, 4, and 5. We associate the aggregate system with its aggregate graph. Then:*

1. The aggregate system is structurally a candidate oscillator in the strong sense iff all the cycles in the aggregate graph are negative.
2. The aggregate system is a candidate multistationary system in the strong sense iff all the cycles in the aggregate graph are positive.

The proof is reported in Appendix F. The MAPK pathway is an excellent example of monotone aggregate system, where each stage of the phosphorylation cascade can be regarded as a monotone module [5]. It is well known that, depending on the active feedback loops, this network can generate bistable or oscillatory behaviors[5, 18, 41].

## 4 Discussion and Conclusions

We described a general, loop–based classification of multistationary and oscillatory behaviors in dynamical systems with sign–definite Jacobian. This classification builds on the well known Thomas' conjectures, and distinguishes between strong and weak candidate oscillators and multistationary systems depending on the presence of exclusively negative or exclusively positive, or coexisting positive and negative loops. We provide a complete characterization, where necessary and sufficient conditions rely on the ability of locally scaling the entries of the Jacobian. We say that this characterization is robust with respect to parametric uncertainty in a model, because it only depends on the sign–definite Jacobian of the model.

Parameter–free criteria to identify the possible dynamic phenotype of a system are particularly important in the context of molecular systems. Many biological network models are sign definite, and can thus be appropriately classified according to our framework; we have mentioned a few notable examples, which include the famous Elowitz repressilator and a variety of toggle switches [6, 16, 38]. Realistic, detailed models for molecular and biochemical networks are most likely to fall into the category of weak candidate oscillator or multistationary systems; however, due to the complexity of the cellular environment, such models may not enjoy the sign definiteness property and thus be outside of the scope of this work.

Other formalisms have been developed in the past to achieve structural conclusions on the behavior of biochemical systems. The chemical reaction network theory provides many structural results; the zero deficiency theorem [17], for instance, rules out the presence of oscillations and multistationary behavior regardless of specific kinetic rates. In the broad context of chemical reaction networks, loop conditions associated with the presence of Hopf or pitchfork instability have been formulated for systems governed by mass–action kinetics using species–reaction graphs [9, 10, 30, 35, 36] and algebraic geometry [13, 14]. While chemical reaction networks are undoubtedly a very important class of models, phenomenological models are often preferred in biology when pathways are not known with sufficient mechanistic detail. Thus, we argue that Jacobian graphical or algebraic conditions may have more general applicability than species–reaction graph conditions [11], which are built upon the mass–action kinetic formalism and cannot account for Hill functions or qualitative relationships.

Another class of very powerful robust analysis methods is given by the theory of monotone systems [44, 46], which enables the simplification of large, complex systems into interconnections of gray–box, input/output monotonic subnetworks [3]. Indeed, monotonicity and existence of steady–state characteristics are properties that facilitate the detection of multistationary behaviors in systems of arbitrary size [5]. Monotonicity, accompanied by small–gain conditions, is used to provide necessary conditions for oscillations in [4]. Because a monotone system within a large network can be collapsed into an element with a sign–definite input/output mapping, we have easily extended our classification to interconnections of monotone subsystems.

Among the limitations of this work, we point out that our results hold for systems in which interactions between nodes are independent. Models built using mass action kinetics, for example, do not fall under this category. For instance, take the reaction 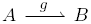 The corresponding concentration dynamics of *A* will include a term −*g(A)*, representing the consumption of *A*; the same term will appear, with opposite sign, in the concentration dynamics of *B*. This situation introduces constraints in the magnitude of the Jacobian entries, which we have not taken into account. A more detailed discussion on this point is reported in Appendix E.

We also remark that we considered only networks that are non–critical, i.e. the length of any cycle in the system structure is larger than two. If this assumption does not hold true, then our characterization of candidate oscillators is not fully valid, as discussed in Appendix C.

We believe that our classification is useful for the *design* of artificial *in vitro* biochemical networks. Recently, *in vitro* reaction environments have gained significant attention because of their reduced and controllable complexity. In particular, synthetic multistationary and oscillatory networks composed of nucleic acids and proteins have been built with systematic design methods [19, 28, 29, 37]; these networks promise to be programmable components in larger, integrated systems for nanomanufacturing, pattern formation, and artificial biomaterials. Although created with a bottom–up approach, designing reaction by reaction, these synthetic *in vitro* networks still suffer from parametric uncertainty and undesired dynamics [19], especially due to the variability in purity and activity of key components. We foresee that by following and enforcing loop sign conditions suggested in this paper, it will be possible to increase the tunability and robustness of these artificial networks.

## Acknowledgments

The authors would like to thank Professor F. Zanolin for his extremely valuable suggestions and J. Kim for his feedback on this manuscript. Elisa Franco acknowledges financial support from NSF grant CMMI–1266402 and from the Bourns College of Engineering at the University of California at Riverside.

## Appendices

### A Nondimensionalization of the two–gene network

We will carry out the nondimensionalization procedure for the toggle switch network, leaving the derivation for the other cases to the reader. We follow nondimensionalization steps similar to those proposed in [16] and [19, 28]. Consider the simple (dimensional) model:

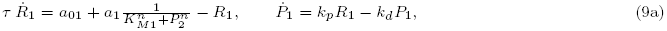

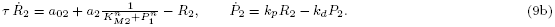

Here a_0i_ is the “leak” transcription of RNA; for simplicity we assume that the translation and degradation rates for the proteins are the same. Constant τ is the mRNA half–life in the system. Constants *K_Mi_* represent the number of proteins necessary to half–maximally repress *R_i_*. Finally, assume the translation efficiency of each RNA species is given by *p̄_i_*, which corresponds to the average number of proteins produced by a single RNA molecule.

We define the nondimensional variables: 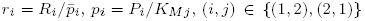. We rescale time as *t̃* = *t/τ*, and also define the nondimensional parameters:

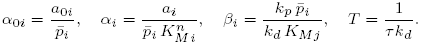

The resulting nondimensional equations are:

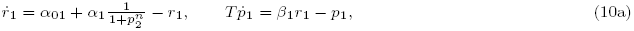

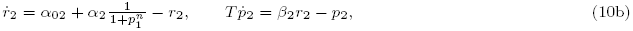

Finally, if we assume *T* ≈ 1, we get a system in the same form as equations (1).

### B Proofs of the main results

In this appendix we report the complete proofs of our main results. We begin by introducing a transformation employed throughout the proofs to find realization of structures satisfying a property of interest. This transformation is particularly useful in the necessity part of the proofs.

*Remark 2* Some of our proofs are built with the following argument. Suppose we are given a condition *𝒞* on a structure associated with a system *ẋ*(*t*) = *f*(*x*(*t*)), and we want to prove that the condition is necessary for a certain property *𝒫*. We reason by contradiction, supposing condition *𝒞* is not satisfied by the structure; then, we show that there always exists a realization of the structure for which *𝒫* fails.

#### B.1 Vector field transformations by *ν_K,∈_*–functions

As a preliminary technical step we introduce the class of *ν_k,∈_*–functions, where *κ* and *∈* > 0 are real parameters.

##### Definition 13

A *ν_K,∈_* function (see Figure 5) is a strictly increasing, continuously differentiable, odd function (i.e. *ν_K,∈_*(*x*) = – *ν_K,∈_*(‒*x*)), satisfying:

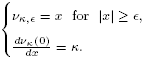

A function of class *ν_K,∈_* has a scalable derivative at the origin and is the identity function outside the *∈*–ball.

**Example** Among the infinite possible *ν_K,∈_*–functions, we consider this example for illustrative purposes:

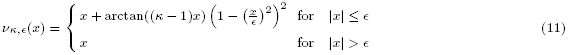

The derivative of function (11) inside the interval [−*∈*, *∈*] is

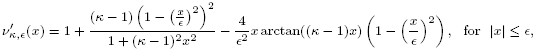

while 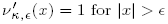. For all *κ* > 0, its derivative 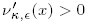 is continuous and positive, and its derivative in *x* = 0 is equal to *κ* as desired. This example function is plotted in Figure 5.

**Fig. 5.**
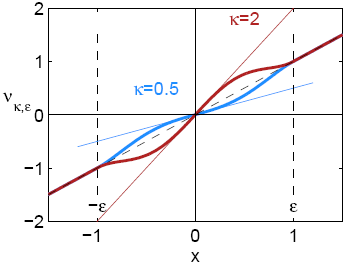
Plot of the example *ν_κ,∈_ (x)*–function in equation (11), with *∈* = 1, *κ* = 2 (red) and *κ* = 0.5 (blue).

**Transformation by** *ν_κ,∈_* **function:** Let us compose the vector field *f(x)* and a function of class *ν_κ,∈_*, i.e. we introduce a transformation:

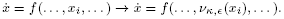

This operation:

a. does not chnge the equilibrium *x_i_* = 0;
b. does not change the structure of the system (does not alter the sign of the Jacoian entries);
c. hanges the partial derivatives in *x_i_* = 0 as:

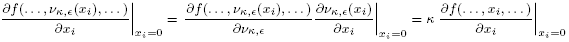
d. *ẋ* = *f(x)* is unchange outsie the ∈-ball after the transformation. In particular, bouneness properties of the solution are not affected.

If we apply this transformation to a vector field *f(x)* in a neighborhood of the origin as an equilibrium point, the elements of the Jacobian of *f* at *x* = 0 can be arbitrarily scaled.

*Remark 3* We are going to use *ν_κ,∈_ (x)* transformations in the following proofs. Given a structure, we will exploit a *ν_κ,∈_ (x)* transformation to find a realization that satisfies the property of interest. An important advantage of *ν_κ,∈_ (x)* transformations is that they preserve the boundedness of the system solution.

#### B.2 Proof of Proposition 1

**Necessity**. If there are no negative cycles, then the system is monotone, and it does not admit oscillatory dynamics (see, for instance, the survey [46]). More precisely, if the system is monotone then its Jacobian is a Metzler matrix, hence it has a dominant real eigenvalue so that any simple transition to instability is an MTI.

**Sufficiency**. The idea of this proof is simple: suppose there exists a single negative loop; then we apply a *ν_k,∈_* transformation to all remaining loops, scaling down the *κ* parameter until we virtually “eliminate” all the other loops. For instance, in the structure shown in Figure 6a we can select the cycle formed by nodes 1-2-3, and consider the new graph achieved by considering only the orange interconnections, Figure 6b.

Then for each arc, or Jacobian entry, we introduce a transformation with a *ν_k,∈_* function, and we can scale the interconnections so that *κ_ij_* > 1 for the arcs involved in the negative loop, while small values *k_ij_* ≪ 1 are assigned to the arcs not involved in the loop. By reordering the nodes, we can always find a realization having a Jacobian (or a principal submatrix of the Jacobian) of the form:

**Fig. 6.**
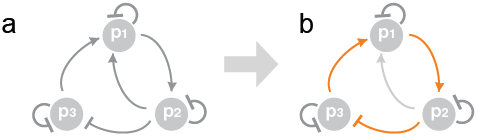
a. Example of incoherent oscillator presented in [38]. b. Emphasis on the negative loop (orange).

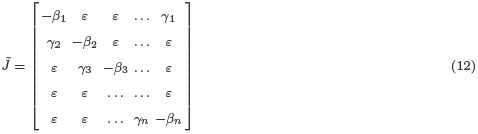

where *β*_i_ > 0, and *γ*_i_ ≠ 0. With an abuse of notation, the entries corresponding to interconnections scaled to arbitrarily small values are denoted as *ε* ≈ 0, instead of individual terms *ε_i,j_*. By assumption

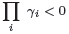

Neglecting the ε–entries, which can be arbitrarily small, the realization Jacobian matrix *J̃* has characteristic polynomial

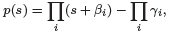

whose coefficients are all positive. Therefore *J* has no real nonnegative eigenvalues.

Elementary root locus techniques show that some of the eigenvalues may become unstable (necessarily complex) for large *γ_i_* (while they remain stable for suitably small *γ_i_*). Therefore, there is always a choice of parameters yielding an OTI.

#### B.3 Proof of Proposition 2

Necessity Assume there is a positive cycle in the structure. We can apply the *ν_k,∈_* transformation, scaling up the positive cycle and scaling down arbitrarily close to zero all the other loops, achieving a realization Jacobian similar to matrix (12). Proceeding as in the sufficiency part of the proof for Proposition 1, we conclude that the characteristic polynomial of the realization is:

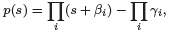

where Π*_i_ γ_i_* > 0 since the loop is positive. Therefore, a realization can be destabilized by driving one real root to cross the imaginary axis through the origin, generating an MTI. (Note the “symmetry” with respect to the sufficiency part of the Π*_i_ γ_i_* < 0.)

**Sufficiency** Assume that the structure (sign pattern matrix associated with the Jacobian) has only negative cycles. The characteristic polynomial corresponding to the structure is:

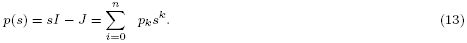

It is well known that the characteristic polynomial is always a monic polynomial, i.e. *p_n_* = 1. In addition:

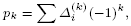

where *Δ_i_^(k)^* are the determinants of all the leading minors of order *k* of the Jacobian. We now invoke the following property, from Theorem 3.1 in [33]:

*Property 1* If all the diagonal entries of *J* are negative and if all the cycles are non–positive, then each leading minor of order *k* has sign (−1)^*k*^.

This property implies that all the coefficients of the characteristic polynomial *p(s)* are positive. In turn, this implies that no real non–negative roots are admitted. Therefore we conclude that if the system is destabilized, then a complex conjugate pair of eigenvalues with nonnegative real part must exist (OTI).

#### B.4 Proof of Corollary 1

The proof of Corollary 1 relies on the so–called degree theory. We assume that the system of interest is a candidate strong oscillator, and therefore its structure has only negative loops. In view of the result proposed in [33], which we just used in the proof of Proposition 2, we have that the determinant of a realization matrix *J(x)* with only negative loops, at *any* equilibrium point *x̄*, satisfies:

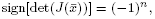

where *n* is the system dimension. Moreover, sign[det(*J(x̄)*)] does not change as a function of *x̄*. Now, let us denote the equilibria as *x̄_i_*. Degree theory also provides us with this equality, which holds for globally bounded flows (see also Lemma 2 in [24], or reference [39]):

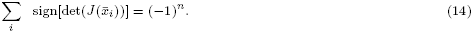

Because in our case all the terms of the sum have the same sign, there must be a single equilibrium. Thus, a strong candidate oscillator structure admits a single equilibrium.

#### B.5 Proof of Proposition 3

**Necessity** If there are no positive cycles, the system is a candidate oscillator in the strong sense. We have seen that the determinant of the Jacobian is never 0, and therefore there cannot be 0 eigenvalues. Thus, no simple MTI can occur.

**Sufficiency** Assume that there exists a positive cycle. Then using the *ν_κ_* function arguments we can prove the existence of a realization with a simple MTI by means of the same argument given in the necessity part of the proof of Proposition 2.

#### B.6 Proof of Proposition 4

**Necessity** If not all the cycles are positive, then there exists a negative cycle. Thus, the system is a candidate oscillator in the weak sense. Then, we can find a realization exhibiting an OTI.

**Sufficiency** If all the cycles are positive, then the system is monotone. Therefore, the Jacobian always has a dominant real eigenvalue. If the system admits a simple transition to instability, it must be monotonic [44, 46].

#### B.7 Proof of Corollary 2

We assume that the system is a candidate multistationary in the strong sense (hence its structure has only positive cycles), therefore any simple transition to instability must be an MTI. This means that, for any realization, the determinant of the Jacobian changes sign when parameter *μ* crosses the critical value *μ*\*. In other words, for values of *μ* to the left of *μ*\* the characteristic polynomial must have a positive constant term, and thus 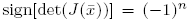. For values of *μ* to the right of *μ*\* we have 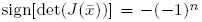. This means that equality (14) cannot be true in a right neighborhood of *μ*\*, unless additional equilibrium points appear. If the additional equilibria are non–singular, i.e. they have a sign–definite Jacobian, then there are at least two of them.

Finally, let us assume that exactly two additional equilibria appear. Assume our original equilibrium is *x̄* = 0, unstable by assumption; let *x_i_, i* = 1, 2 be additional equilibria introduced by the transition to instability. In a right neighborhood of *μ*\*, consider the two regions

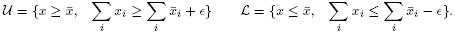

In view of the assumptions of a) monotonicity of the system, b) instability of *x̄* and c) irreducibility of the Jacobian, both regions *𝒰* and *ℒ* are positively invariant. Given our assumption of boundedness of the system solution, *𝒰* and *ℒ* are both bounded. Then, each set must include one of the two equilibrium points^2^.

To conclude the proof, we invoke a known result in the context of monotone systems (see, for instance [26], page 458 Theorem D): since the two equilibria are unique in the bounded sets *𝒰* and *ℒ*, they must be stable. The two–gene positive feedback system and the toggle switch are clear examples of this result.

### C Critical systems

Here we discuss our results in the context of critical systems. Starting from system (3) and the related assumptions listed in Section 2, we provide the following definition:

#### Definition 14

System (3) is critical iff all the negative cycles of its structure (if any) are of order two.

We now examine the validity of our characterization for critical systems.

*Validity of Proposition 1:* For critical networks, the sufficiency part of Proposition 1 does not hold: the existence of negative cycles of length 2 does not assure that the system can oscillate. For example, consider the following system:

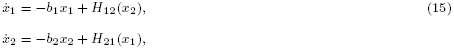

where *b*_1_ and *b*_2_ are positive constants, and *H_ij_* are positive, bounded, and monotonic functions; to ensure the existence of a negative loop, we assume that (for example) *H*_12_ is non–increasing, while *H*_21_ is non–decreasing. (We can take *H_ij_* as Hill functions). For any choice of *b_i_* and *H_ij_*, the system has a single, asymptotically stable equilibrium point. However, oscillations are possible if we take b_1_ = b_2_ = 0, i.e. if we remove our assumption that negative self loops must exist at each node.

The necessity part of Proposition 1 still holds, because if there are no negative cycles the system cannot have equilibria with oscillatory instability.

*Validity of Proposition 2:* In the presence of negative cycles only, then instability must be oscillatory; thus, Proposition 2 still holds. However, some critical systems cannot be destabilized at all, as just shown in example (15).

It is legitimate to ask whether a critical system with only negative loops (all of order two) is necessarily stable. This would extend Proposition 1 as follows:

*Conjecture:* A system is a candidate oscillator in the weak sense if and only if it has at least a negative cycle of order greater than 2.

While the “if” part of this conjecture is true, unfortunately the the “'only if” part is false. This can be seen with a counterexample: we modify example (15) by introducing a new variable *x*_3_ and a positive loop:

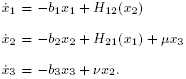

The system Jacobian is:

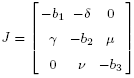

This matrix may well have unstable complex eigenvalues. Let us assume *b*_1_ = *b*_2_ = 0. The characteristic polynomial is:

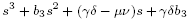

Building the Routh–Hurwitz table for this polynomial, one can check that for *μ*, *ν* > 0 two roots are complex with positive real parts. For b_1_,b_2_ > 0 small enough, the property is preserved.

A property relevant to our discussion on critical cases is the following known result (see for instance [15] Chapter 6.5):

#### Proposition 6

*If all the self–loops are negative, and all the other loops are negative and of order two, then any equilibrium of the system is stable*.

### D Systems affected by delays

Many molecular systems have been successfully modeled using delayed differential equations. A notable example is given by gene networks, where the transport of RNA and proteins across cellular membranes are well captured by explicit delays [25, 31]. Delay differential equations are infinite dimensional systems and a formal, exhaustive treatment of this case would require a more sophisticated setup. However, we can show that local, loop–based sufficient conditions for OTIs and MTIs can be stated in a wide class of systems with delays. Consider the delay differential equation model:

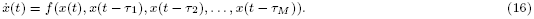

Under standard differentiability assumptions, the corresponding linearized system around an equilibrium point is:

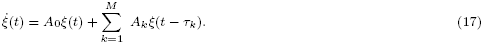

For simplicity we assume that there are no delayed self–loops, which means that the matrices *A_k_* have zero diagonal entries for *k* ≥ 1.

It is well known that the stability of the system above can be established by inspecting the roots of this equation:

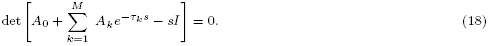

Stability is ensured if the roots have negative real parts. We now consider the following auxiliary system, in which all delays are (fictitiously) set to 0:

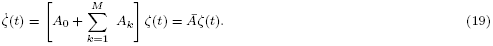

Matrix *Ā* is the same Jacobian we would obtain by setting the delays *τ_i_* = 0 in system (16):

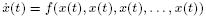

Note that equilibria are delay–independent, so the delay–free system above has the same equilibria as the delayed system (16).

As previously done in this paper, we can associate graphs with systems (17) and (19), where nodes correspond to species and signed, directed arcs correspond to the dynamic interactions among species (delayed or not), defined by matrices *A_k_, k* = 0,..., *M*. Delays do not change the sign of the loops. Therefore any positive/negative cycle of (17) corresponds to a positive/negative cycle of (19).

#### Proposition 7

*Assume system* (17) *has only negative loops. Then, the system admits solely OTIs*.

**Proof** *Ab absurdo*, assume that the system admits a monotonic transition to instability, and thus one root of equation (18) crosses the imaginary axis with value *s* = 0. This is equivalent to writing equation (18) as:

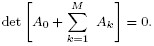

If this is true, then also the auxiliary system (19) must admit a zero eigenvalue. But this is impossible if the system has only negative loops, according to Proposition 2.

#### Proposition 8

*Assume that system* (17) *has only positive loops. Then, the system admits solely MTIs*.

**Proof** To prove this proposition, we need two observations:

a. In the bsence of negtive loops (with the exception of self–loops), system (17) is liner positive system with dely. Therefore, mtrix *Ā* in system (19) is Metzler mtrix.
b. We invoke a well known property of *positive* linear systems with delay (with no delayed self-loops). System (18) is stable if and only if its delay–free, auxiliary counterpart (19) is stable [23, 32] (see also [52], Theorem 6.5).

If all its loops are positive, the auxiliary system (19) can transition to instability only by means of a pole in *s* = 0.Again *ab absurdo* let us assume that the delay system (18) admits instability with a pair of dominant imaginary eigenvalues and no real eigenvalue crossing the origin. Then, the auxiliary system (19) would also be unstable. However, the dominant eigenvalue of the Metzler matrix *Ā* is real; therefore the auxiliary system (19) would transition to instability with a pole at *s* = 0. But this would also imply that equation (18) is satisfied with *s* = 0, which is a contradiction. Therefore, we conclude that if all its loops are positive, a transition to instability with a pair of imaginary eigenvalues is impossible for system (17).

As a comment to the above proposition, we stress that equilibria are delay–independent, therefore if the auxiliary subsystem presents multiple equilibria, so does the delayed system.

We conclude this appendix by noting that Proposition 6 is not valid in the presence of delays. Indeed, the presence of a delay in a “second order negative loop” may compromise stability, as it is well known in control theory.

### E Structural cross–constraints among function

We have not considered models presenting cross–linked dynamic terms in several equations. Cross–linked terms appear typically in models built using the mass action kinetics formalism. For instance consider the reactions:

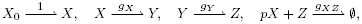

where ∅ indicates elimination of a species from the system (degradation or out–flow). Indicating the concentration of a species with the corresponding small letter, the differential equation model is:

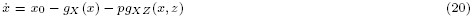

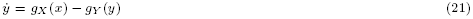

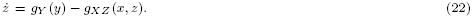

Here, all the reaction rates are increasing functions. Identical dynamic terms appear in the three equations, so the Jacobian has dependent entries:

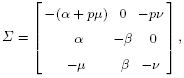

where, for a fixed equilibrium, all Greek letters represent positive constants. This structure presents both negative and positive cycles, but it may only undergo oscillatory transitions to instability; thus, it is a candidate oscillator in the strong sense. This fact can be seen by writing the characteristic polynomial:

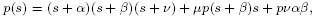

where all coefficients are positive, and thus there cannot be real positive roots.

This would seem in contradiction with the presence of the positive cycle 1 → 2 → 1. However, our results are not invalidated because we do not consider cross–constraints among entries in the Jacobian, such as the fact that *J*_22_ = −*J*_32_. Clearly, if we could change all entries independently (without changing sign) this system could present a monotonic transition to instability.

For systems with cross–constrained dynamics, we believe that algorithmic/numerical methods are the best approach to discriminate admissible transitions to instability.

### F Proof of Proposition 5

We recall that each subsystem composing a monotonic aggregate is described by the following equations:

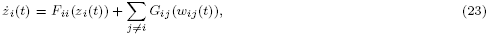

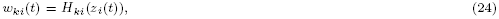

where *z_i_(t)* is the state vector, *w_ij_* are inputs, and *w_ki_* are outputs of the *i*–th subsystem. We assume that each subsystem is unconditionally stable, and that Assumptions 3 and 4 hold.

Unconditional stability implies that for *constant* input values *w̄_ij_*, each subsystem admits a single equilibrium *z̄_i_* defined implicitly by:

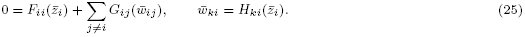

The equilibrium *z̄_i_* is globally stable, i.e. all eigenvalues of the Jacobian *J_i_* have a negative real part. Given the steady state condition (25), we define:

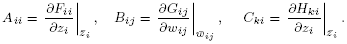

We recall that the system can be associated with an **aggregate graph**, where subsystems are collapsed into nodes, interconnected through directed arcs whose sign depends on the trend (non–decreasing or non–increasing) of function *H_ki_(z_i_)*: thus, the arc sign depends on the sign of *C_ki_*.

#### Lemma 1

*The i–th steady state input–to–output mapping, or steady–state characteristic, is monotonic. The input–to–output mapping between each pair w_ij_, w_ki_ is implicitly defined by equality* (25), *and we find:*

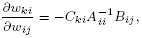

*which is a positive or negative scalar depending on the sign of the elements of C_ki_*.

**Proof** As a consequence of our monotonicity and unconditional stability assumptions Assumption 5), *A*_*ii*_ is a Metzler matrix, and it is asymptotically stable. Therefore, all elements of its inverse *A_*ij*_^−1^*; are negative. By construction Assumption 3) *B_ij_* has non–negative elements, and *C_ki_* has all non–negative elements or all non–positive elements depending on the type of interaction (Assumption 4). Thus, the sign of *∂w_ki_/∂w_ij_* only depends on the sign of *C_ki_*, which concludes our proof.

#### Proof of Proposition 5

*Part 1. Aggregates of monotone systems that are structural candidate oscillators*.

**Necessity** The existence of a negative cycle in the aggregate graph implies the existence of a negative cycle in the standard graph. Thus, this part can be immediately concluded.

**Sufficiency** To prove sufficiency, it is enough to show that the Jacobian of the system cannot have real non–negative eigenvalues. This is equivalent to saying that only oscillatory destabilization is possible. Assume *A* is the Jacobian matrix of the aggregate system (i.e. of the overall interconnection of monotonic subsystems). We will show that:

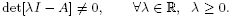

We develop our proof *ab absurdo*. Assume that *A* admits a real non–negative eigenvalue *λ* ≥ 0. Denote the *i*–th linearized subsystem as:

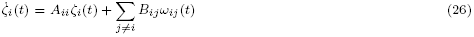

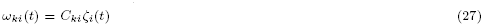

Therefore, any eigenvalue *λ* must satisfy the equation:

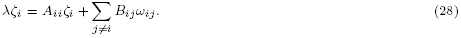

We find:

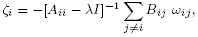

where all the elements of [*A_ii_–λI*]^−1^ are negative. (This is a consequence of monotonicity: *A*_*ii*_ is a stable Metzler matrix; [*A_ii_–λI*] is still a stable Metzler matrix, because *λ* ≥ 0; thus all the elements of [*A_ii_–λI*]^−1^ are negative.)

Then, we can write:

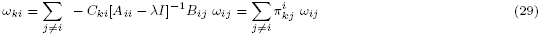

where 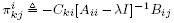 are scalars. Equations (29) are linear in *ω*_*ki*_, and we can rewrite them in a compact form

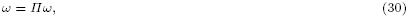

where *ω* is a vector including all the interconnection variables *ω_ij_*. Variables *ω_ij_* can be considered “arc–variables”, because they define the interconnections in the aggregate system. Therefore, the sign of the aggregate graph arc from node *i* to node *k* depends on the sign of 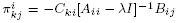. Thus, matrix *Π* appearing in equation (30) is characterized by the same cycles defined by the Jacobian *A* of the aggregate system. Let *Σ_Π_* be the sign matrix corresponding to *Π*:

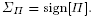

Every cycle in the aggregate graph corresponds to a cycle in matrix *Σ_Π_*. For example, the sign matrix *Σ_Π_* associated with the aggregate graph in Figure 7 is a 5 × 5 matrix (since there are 5 “arcs”); if we order the arc variables in vector *ω* = [*ω*_21_ *ω*_32_ *ω*_43_ *ω*_14_ *ω*_32_], we find matrix *Σ_Π_* in expression 31.

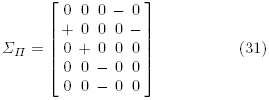

**Fig. 7.**
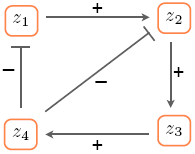
Aggregate graph, corresponding to the sign matrix (31).

Since any cycle of the graph corresponds to a cycle of *Σ_Π_* (as evidenced in the above example), if all the cycles in the aggregate graph are negative, then also all the cycles in matrix *Σ_Π_* are negative. As a consequence, the determinant of matrix [*Π – I*] is positive. Now we go back to relation (30), which is equivalent to [*Π – I*]*ω* = 0: this relation implies det[*Π – I*] = 0. However, we encounter a contradiction: if all the cycles are negative, we just showed that det[*Π – I*] > 0, thus [*Π – I*]*ω* = 0 cannot be true for *ω* ≠ 0. Therefore, we conclude that the system does not admit non–negative eigenvalues *λ* ≥ 0.

*Part 2. Aggregates of monotone systems that are structural candidate multistationary systems*.

Necessity and sufficiency can be built exactly as done for the proof provided for aggregates of monotone systems that are structural oscillatory candidates. Thus, we omit these demonstrations.

## Footnotes

1 This definition holds for systems transitioning to instability from the right to the left neighborhood of *μ*\* in the above definition, it suffices to take *μ̂* = *μ*\* – *μ* as the bifurcation parameter.

2 Exploiting the Jacobian irreducibility assumption, it is actually possible to prove that *at least* two additional equilibria arise.

